# Floss-Mediated Gingival Mucosal Immunization with HBc-E18-3 VLPs Induces Long-Lasting Intestinal IgG and Provides a Candidate Strategy for Intervention of FcRn-Related Autoimmune Injury

**DOI:** 10.64898/2026.08.10.743934

**Authors:** Tingting Zhai, Shuai Jiang

## Abstract

Echovirus 18 (E18) is a predominant pathogen causing aseptic meningitis in children, and post-E18 infection frequently triggers myasthenia gravis-like autoimmune neurological damage. This pathological process relies on neonatal Fc receptor (FcRn)-mediated IgG transcytosis across mucosal barriers, and FcRn also acts as an essential functional receptor required for E18 attachment and uncoating during host cell invasion. At present, no E18-specific prophylactic vaccine has been clinically approved, and anti-FcRn monoclonal antibodies are the available therapeutics to alleviate autoantibody-mediated tissue injury.

We constructed an integrated automated phylogenetic pipeline named *evolution_conservation*, which enables rapid tracing of the evolutionary position and genetic relatedness of clinical isolates to identify closely related strains from previous outbreaks. Serving as an in silico alternative to animal experiments, this pipeline supports reference-guided vaccine design and longitudinal comparative assessment of vaccine safety and efficacy, facilitates identification of patient populations presenting rare post-viral sequelae, and accelerates clinical trial progression. In this study, we inserted the pre-screened linear epitope E18-3 into a truncated hepatitis B core (HBc) scaffold to generate chimeric virus-like particles (VLPs). A non-invasive floss-based gingival mucosal immunization mouse model was established, with subcutaneous Freund’s adjuvant immunization set as the control group. ELISA results confirmed that gingival mucosal delivery of particulate HBc-E18-3 VLPs alone could induce sustained high levels of antigen-specific intestinal IgG *in vivo*.

Drawing on research paradigms of therapeutic neoantigen vaccines for tumor recurrence prevention, the *evolution_conservation* bioinformatic pipeline and mucosal VLP platform described herein establish an innovative framework for developing antigen-competitive prophylactic and therapeutic vaccines targeting FcRn for myasthenia gravis and autoimmune encephalitis.

## 1 Introduction

Echovirus 18 (E18), a member of Enterovirus B, is a non-enveloped single-stranded RNA virus with a spherical shape (∼25 nm diameter) exhibiting icosahedral symmetry. Humans are its primary natural host, and viral transmission occurs mainly via the fecal–oral route. After replicating in intestinal epithelial cells, E18 invades the central nervous system and induces febrile aseptic meningitis in children^1–7^. Echoviruses were first isolated from patient feces during the 1951 poliovirus epidemic as non-polio enteroviruses capable of triggering cytopathic effects in cultured cells. The paralytic sequelae of poliovirus infection share clinical similarities with myasthenia gravis—the primary indication of clinically approved anti-FcRn monoclonal antibodies^8,9^—and FcRn functions as a dual receptor mediating both E18 attachment and capsid uncoating^10,11^. To date, no dedicated prophylactic vaccine against E18 has received clinical authorization. Approved anti-FcRn agents such as efgartigimod and rozanolixizumab compete with pathogenic IgG for FcRn binding to reduce IgG recycling efficiency and circulating antibody concentrations^12–14^, yet these drugs fail to block primary viral infection or prevent recurrent episodes of autoimmune diseases including myasthenia gravis and autoimmune encephalitis.

Clinical studies have demonstrated that the gut microbiome of rheumatoid arthritis (RA) patients accumulates specific bacteriophages (e.g., *Prevotella* and *Oscillibacter* phages), whose peptides share high sequence homology with the RA autoantigen BiP. Via molecular mimicry, these phage-derived epitopes activate CD4^⁺^ T cells and plasma cells, initiating autoimmune responses and driving RA progression^15^. Virus-like particles (VLPs) self-assemble from recombinant viral structural proteins without replicative genomes; they faithfully recapitulate native antigen conformations, robustly activate B and T lymphocytes, and carry no infectious risk. Truncated HBc (amino acids 1–149) self-assembles into homogeneous T=4 nanoparticles, and the surface-exposed loop spanning residues 78–79 represents the optimal insertion site for exogenous linear epitopes^16–21^. CpG oligodeoxynucleotides are clinically approved human vaccine TLR9 agonists that synergistically boost systemic and mucosal humoral immune responses^22–24^. The junctional epithelium beneath the gingival sulcus exhibits high antigen permeability and is enriched with innate immune cells^25,26^; topical administration via antigen-coated dental floss enables non-invasive mucosal immunization^27,28^.

Owing to interspecies physiological disparities between humans and laboratory animals^29^, conventional animal models for biologic safety and efficacy evaluation rarely translate to successful clinical outcomes, accounting for the ∼90% failure rate of novel drug candidates^30,31^. Humans, particularly children, constitute the natural susceptible host for E18, while mice are non-permissive to E18 infection, rendering traditional viral challenge and autoimmune animal models ineffective for predictive preclinical assessment. Against this backdrop, three critical research gaps were identified:

1. How to develop computational alternatives to animal assays to accelerate research efficiency and improve clinical translatability?
2. Floss-based vaccination improves vaccine accessibility, yet inactivated whole-virus vaccines carry biosafety risks and isolated linear peptide antigens display extremely weak intrinsic immunogenicity. It remains unclear how HBc-VLPs simultaneously presenting HBc conformational epitopes and viral linear epitopes elicit protective neutralizing antibodies following floss-mediated mucosal immunization.
3. Existing HBc-VLP research predominantly focuses on prophylactic antiviral applications, with minimal exploration of their potential to modulate autoimmune disease progression via the IgG-FcRn pathological axis; few design frameworks extend HBc-VLPs as autoimmune therapeutic vaccines.

To address these limitations, we developed an integrated Python-based bioinformatic pipeline *evolution_conservation*, which streamlines batch sequence retrieval, quality control filtering, high-accuracy multiple sequence alignment, quantitative conservation analysis, mutation statistics, and phylogenetic tree visualization within a single executable script. The pipeline eliminates repeated software switching and file import/export, rapidly locates unknown clinical isolates within global echovirus phylogenetic clades, efficiently evaluates genetic relatedness of emerging variant strains, simplifies molecular epidemiological workflows for enteroviruses, and drastically increases analytical throughput. Based on the pre-validated E18-3 epitope^32^, we constructed chimeric HBc-E18-3 VLPs and established a non-invasive floss-mediated gingival mucosal immunization mouse model, with subcutaneous Freund’s adjuvant immunization serving as the systemic immunization control. Integrating documented FcRn molecular mechanisms and the clinical landscape of anti-FcRn monoclonal antibodies, we draw parallels with neoantigen therapeutic vaccine research for oncology and explore the dual translational potential of this mucosal VLP platform for antiviral prophylaxis and FcRn-associated autoimmune disease intervention.

## 2 Materials and Methods

### 2.1 Automated E18 VP1 Bioinformatic Analysis Using the In-House *evolution_conservation* Pipeline

The E18-3 antigen fragment utilized for vaccine construction was identified via systematic epitope screening performed previously in our laboratory; full sequence characteristics and pre-experimental variation datasets are provided in supplementary materials. The integrated automated pipeline *evolution_conservation* consolidates batch sequence retrieval, quality filtering, high-precision MAFFT alignment, neighbor-joining (NJ) phylogenetic tree construction, and visualization workflows. The entire analytical workflow operates via a single script without manual switching between independent software suites, enabling rapid localization of clinical isolates within global echovirus phylogenetic branches and streamlined assessment of genetic relatedness among emerging variants, substantially improving the efficiency of enterovirus molecular epidemiological analysis. All visualized outputs generated by the pipeline are presented in Figure 1.

**Figure 1.**
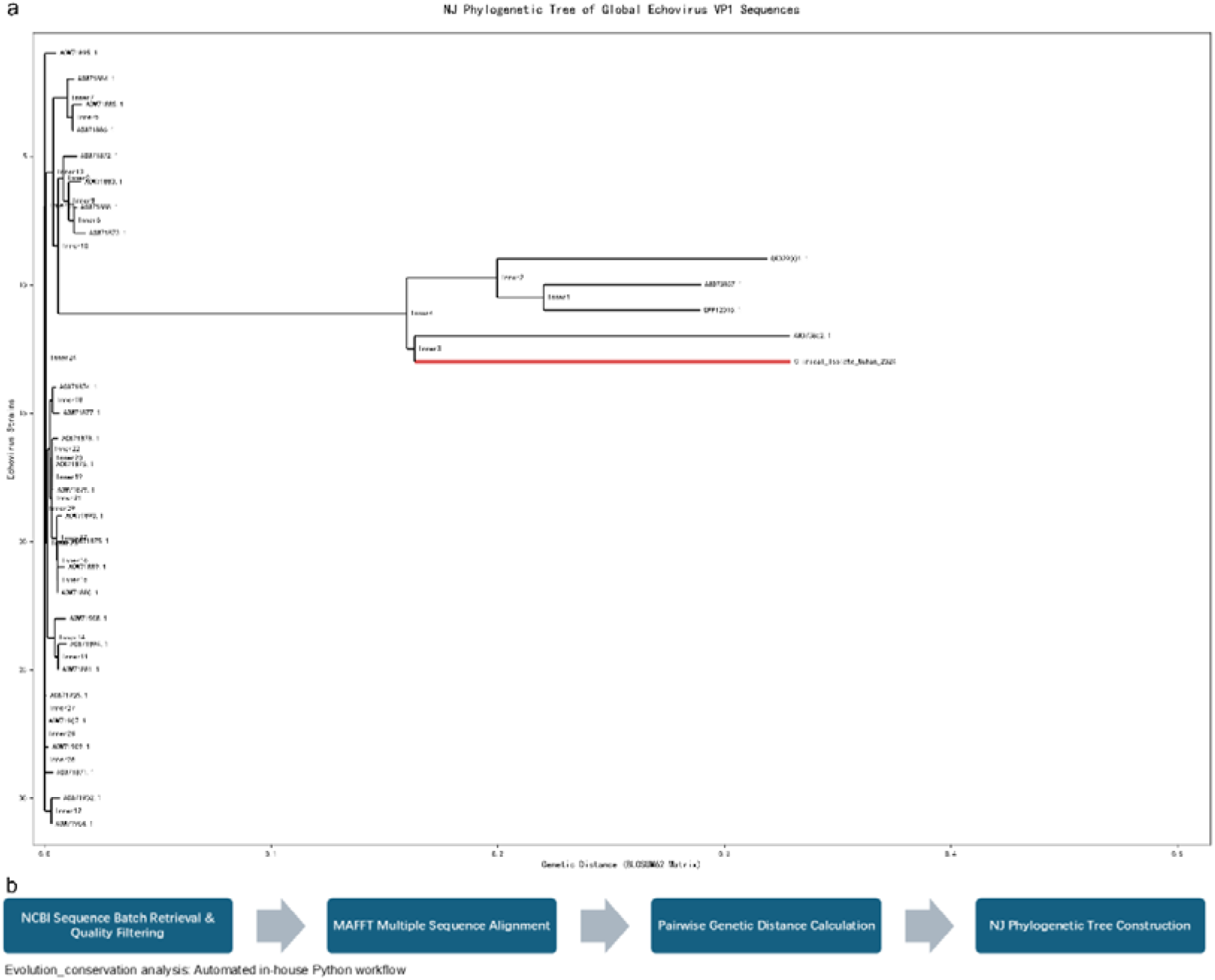
Schematic overview of global E18 VP1 evolutionary analysis using the in-house *evolution_conservation* bioinformatic pipeline. (a) Neighbor-joining (NJ) phylogenetic tree of global E18 VP1 clinical isolates. The clade corresponding to the local Wuhan clinical isolate is highlighted with bold red branches to visualize its genetic relatedness to globally circulating strains. (b) Workflow schematic of the integrated Python pipeline *evolution_conservation*, including batch sequence retrieval, multiple sequence alignment, quantitative conservation analysis, mutation statistics, phylogenetic tree construction, visualization, and bulk export of quantitative statistical tables.

#### 2.1.1 Sequence Retrieval and Quality Control

Clinical Echovirus VP1 sequences were batch-downloaded from NCBI GenBank using the search query: Echovirus VP1 partial clinical isolate NOT polyprotein, with a default maximum batch download limit of 120 sequences (parameter adjustable). Full-length VP1 amino acid sequences derived from an E18 clinical isolate collected from a pediatric meningitis patient in Wuhan were integrated to build a comprehensive sequence library. All FASTA sequences underwent preliminary screening based on amino acid length (280–300 aa), and redundant duplicate sequences were eliminated via MD5 hash calculation of translated amino acid sequences.

#### 2.1.2 Multiple Sequence Alignment

High-accuracy multiple sequence alignment was performed using MAFFT with core parameters: --auto --adjustdirectionaccurately --op 1.0 --ep 0.1. Post-alignment standard FASTA files were exported for subsequent quantitative conservation profiling, mutation counting, and phylogenetic tree reconstruction.

#### 2.1.3 Neighbor-Joining (NJ) Phylogenetic Tree Construction

Pairwise genetic distance matrices were calculated from aligned VP1 sequences using the BLOSUM62 amino acid substitution matrix, and NJ phylogenetic trees were constructed via the Biopython evolutionary module. The clade corresponding to the local Wuhan E18 clinical isolate was highlighted with bold red branches to intuitively illustrate its genetic proximity to globally circulating strains. Phylogenetic trees were exported as high-resolution 300 dpi images.

### 2.2 Construction, Expression and Purification of Recombinant HBc-E18-3 Plasmid

The amino acid sequence of the pre-screened E18-3 epitope is as follows: PVLTHQIMYVPPGGPIPAKVDSYEWQTSTNPSVFWTE. The E18-3 peptide was inserted between residues 78 and 79 of truncated HBc (1–149 aa) via a flexible GGG linker, using the pET-28a (+) vector with a C-terminal 6×His tag retained (Figure 2a, 2b). The codon-optimized recombinant plasmid for *Escherichia coli* expression was synthesized by GenScript Biotech Corporation.

**Figure 2.**
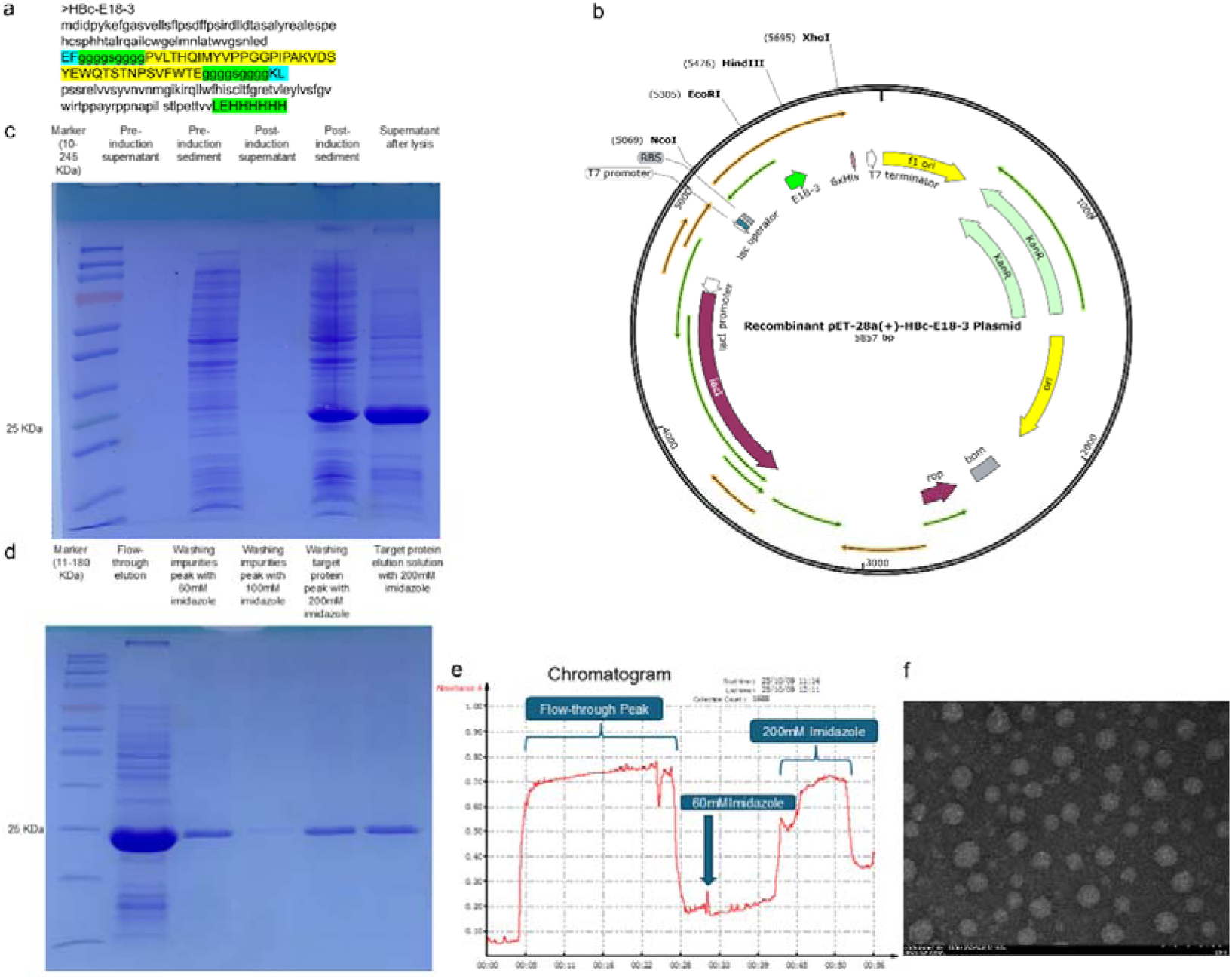
Construction, purification and *in vitro* assembly characterization of recombinant HBc-E18-3 VLPs. (a) Amino acid sequence of chimeric HBc-E18-3. The E18-3 epitope was inserted between residues 78 and 79 of HBc via a flexible GGG linker, with a C-terminal 6×His tag retained. (b) Circular map of the recombinant pET-28a (+)-HBc-E18-3 plasmid (total length: 5857 bp), with key genetic elements annotated including the T7 promoter, NcoI/XhoI restriction enzyme cleavage sites, E18-3 insert fragment, 6×His tag, and kanamycin resistance cassette. (c) SDS-PAGE analysis of whole bacterial lysates before and after IPTG induction of HBc-E18-3. (d) SDS-PAGE profiling of elution fractions collected during denaturing nickel affinity chromatography. (f) Transmission electron micrograph of HBc-E18-3 VLPs self-assembled via gradient urea dialysis, showing homogeneous spherical nanoparticles of ∼30 nm diameter with embedded scale bar.

Recombinant pET-28a (+)-HBc-E18-3 plasmids were transformed into BL21 (DE3) competent cells. Kanamycin-resistant single colonies were cultured in shake flasks at 37 °C, 200 r/min until OD_₆₀₀_ reached 0.6–0.8, followed by induction with 0.5 mM IPTG at 16 °C for 16 h. Cell pellets were harvested by centrifugation at 8000 r/min, 4 °C for 15 min. HBc-E18-3 was expressed as inclusion bodies and purified via nickel affinity chromatography under denaturing conditions (8 M urea) with stepwise imidazole gradient elution (Figure 2c–e). Empty HBc control protein was solubly expressed, with all other expression and purification procedures identical to the chimeric construct.

### 2.3 In Vitro VLP Self-Assembly and Transmission Electron Microscopy (TEM) Characterization

Purified denatured HBc-E18-3 protein was loaded into dialysis bags for stepwise urea removal via gradient dialysis (6 M → 3 M → 1 M → 0 M urea), with each dialysis step incubated at 4 °C for ≥4 h to facilitate spontaneous nanoparticle assembly. Post-assembly supernatants were collected after centrifugation at 12000 r/min, 4 °C for 20 min; protein concentrations were quantified via the BCA assay, and aliquots were stored at −20 °C for subsequent use.

TEM sample preparation utilized phosphotungstic acid negative staining: 10 μL protein solution was applied to carbon-coated copper grids, incubated for 8 min before excess liquid was blotted away, stained with 1% phosphotungstic acid for 5 min, air-dried under an infrared lamp, and imaged to assess nanoparticle size and homogeneity (Figure 2f).

### 2.4 Preparation of Floss-Based Mucosal Vaccine and Subcutaneous Vaccine

#### Floss-delivered vaccine

Purified HBc or HBc-E18-3 protein was thoroughly mixed with an equal mass of CpG adjuvant, evenly coated onto medical dental floss, air-dried in a biosafety cabinet, and stored protected from light at −20 °C. Each floss strip carried a total antigen dose of 25 μg. For gingival immunization, antigen-coated floss was rubbed back and forth 30 times within the gingival sulcus of mouse mandibular incisors to complete mucosal delivery.

#### Freund’s adjuvant subcutaneous vaccine

Each mouse received 100 μg HBc-E18-3 protein emulsified with an equal volume of adjuvant. Complete Freund’s adjuvant was used for primary immunization, and incomplete Freund’s adjuvant for two booster immunizations, administered via multi-site subcutaneous injection at the nape and inguinal regions.

### 2.5 Mouse Immunization Regimen and Sample Collection

Five-week-old female C57BL/6JNifdc mice were purchased from Zhejiang Vital River Laboratory Animal Technology Co., Ltd. Animals were randomly assigned to three experimental groups (n=6 per group): HBc-CpG floss group, HBc-E18-3-CpG floss group, and HBc-E18-3 Freund’s adjuvant subcutaneous group. Immunizations were performed every 14 days, with three total doses administered on Day 0, Day 14, and Day 28.

Fecal, saliva, and serum samples were collected on Day 0, 7, 21, 35, and 56 for quantitative detection of antigen-specific antibodies via ELISA. All mice were sacrificed on Day 65; heart, liver, spleen, lung, and kidney tissues were harvested and fixed for H&E pathological staining to evaluate systemic vaccine safety. All animal procedures were approved by the Animal Ethics Committee of China Pharmaceutical University and strictly performed in accordance with the internationally recognized 3R principles for laboratory animal welfare.

### 2.6 ELISA Detection of Antigen-Specific IgG and IgA

E18-3 peptide, HBc protein, and HBc-E18-3 VLPs were diluted to 100 μg/mL in coating buffer, and 100 μL per well was incubated at 4 °C overnight for plate coating. Plates were washed three times with PBST (PBS supplemented with 0.1% Tween-20), followed by blocking with blocking buffer at 37 °C for 2 h. Serially diluted serum, fecal supernatant, and saliva samples were added as primary antibodies and incubated at 37 °C for 2 h. HRP-conjugated goat anti-mouse IgG/IgA secondary antibodies were diluted 1:5000 and incubated protected from light at 37 °C for 1 h. TMB chromogenic substrate was added, and color development proceeded protected from light at 37 °C for 30 min before termination; absorbance at OD_₄₅₀_ was measured. All data are presented as mean ± standard deviation (n=3 biological replicates per group). Intergroup statistical comparisons were performed using a non-parametric two-tailed Kruskal – Wallis test followed by Dunn’s post-hoc test, with P < 0.05 defined as statistically significant. All results shown are representative of two independent replicate experiments.

### 2.7 Hematoxylin and Eosin (H&E) Staining of Visceral Organs

Heart, liver, spleen, lung, and kidney tissues were dissected from sacrificed mice and fixed in 4% paraformaldehyde. Subsequent paraffin embedding, sectioning, and standard H&E staining were outsourced to Nanjing Tainuowei Biotechnology Co., Ltd. Pathological sections were visualized under an optical microscope to evaluate tissue integrity, inflammatory infiltration, cellular degeneration and necrosis, congestion, and fibrosis for comprehensive assessment of systemic visceral toxicity induced by vaccination.

## 3 Results

### 3.1 Global E18 VP1 Evolutionary Profiling Using the In-House *evolution_conservation* Pipeline

The integrated Python-based bioinformatic pipeline *evolution_conservation* developed in this study enables full-spectrum molecular evolutionary analysis of global E18 VP1 sequences, with all visualized outputs consolidated in Figure 1. This pipeline exhibits two core functional applications:

1. Evolutionary tracing of clinical strains: The pipeline rapidly locates the phylogenetic position of the local Wuhan E18 clinical isolate, which forms a distinct monophyletic clade with a meningitis-derived Vietnamese strain (GenBank accession: AWD73852.1) with minimal genetic distance, confirming sustained co-circulation of E18 strains across the China–Vietnam border.
2. General-purpose epidemiological analytical tool: The pipeline integrates standalone software modules for sequence retrieval, alignment, statistical analysis, and figure generation without manual multi-step processing. By inputting target sequences or search keywords, researchers can instantly assess genetic relatedness of unknown emerging enterovirus clinical isolates, drastically shortening the monitoring cycle for molecular epidemiological surveillance.

The images was exported at a resolution of 300 dpi; supporting statistics for full-length VP1 residues are provided in Supplementary Tables S1 and S2, which corroborate the visualized phylogenetic data in this figure.

### 3.2 Construction and Characterization of Chimeric HBc-E18-3 VLPs

Schematic amino acid sequence of recombinant HBc-E18-3 and circular map of the pET-28a (+) expression plasmid are shown in Figure 2a and 2b. Following IPTG induction, SDS-PAGE detected a specific protein band at ∼24 kDa consistent with the theoretical molecular weight (Figure 2c). Denaturing nickel affinity chromatography yielded a single purified protein band free of contaminating host proteins (Figure 2d, 2e). After urea gradient dialysis to drive *in vitro* self-assembly, transmission electron microscopy revealed homogeneous spherical nanoparticles with a diameter of ∼30 nm, matching the canonical morphology of T=4 HBc VLPs (Figure 2f).

### 3.3 Floss-Mediated Gingival Immunization Induces Sustained Mucosal E18-3-Specific IgG

Schematic gingival mucosal delivery and full vaccination timeline are presented in Figure 3a and 3b. ELISA data collected on Day 35 demonstrated that subcutaneous Freund’s adjuvant immunization induced significantly higher titers of E18-3 linear peptide-specific IgG in feces and saliva relative to floss-based mucosal vaccination; gingival delivery of HBc-E18-3 VLPs failed to elicit detectable neutralizing antibodies targeting isolated linear E18-3 peptide antigen (Figure 3c). Robust high-titer HBc conformational epitope-specific IgG was detected in serum, feces, and saliva from both floss and subcutaneous immunization groups (Figure 3d), verifying that intact particulate HBc nanoparticle architecture is an essential prerequisite for efficient uptake by gingival epithelial antigen-presenting cells and initiation of mucosal adaptive immune responses.

**Figure 3.**
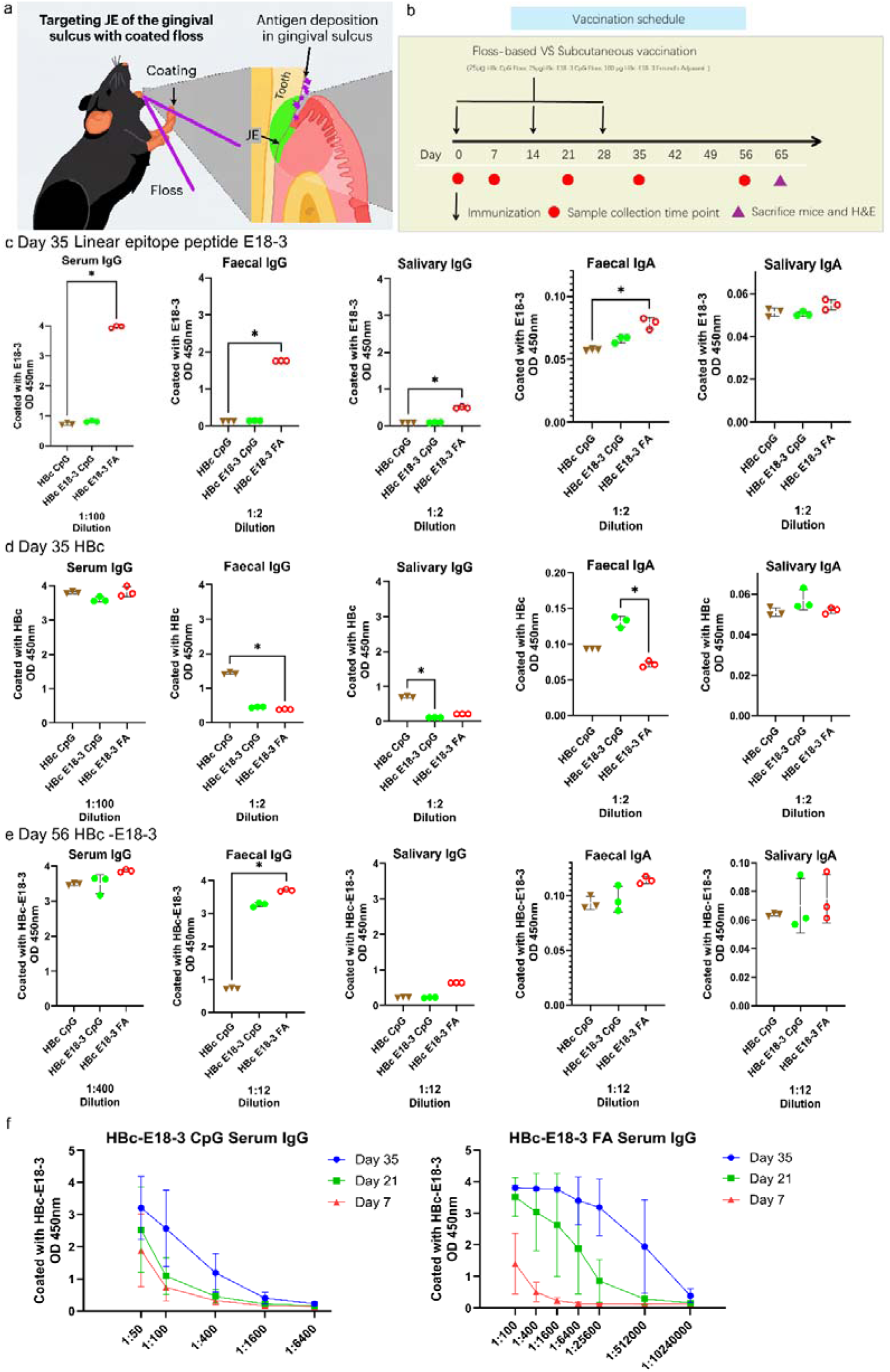
Gingival mucosal vaccination regimen and antigen-specific antibody responses in mice. (a) Schematic illustration of antigen delivery targeting the junctional epithelium (JE) within the mouse gingival sulcus using coated dental floss^27,33^. (b) Complete immunization and sample collection timeline: three immunization doses administered on Day 0, 14, and 28; fecal, saliva, and serum samples collected on Day 7, 21, 35, and 56; all mice sacrificed on Day 65 for pathological tissue analysis. (c) E18-3-specific IgG and IgA levels in feces and saliva measured by ELISA on Day 35 post-immunization. (d) HBc-specific IgG and IgA concentrations in serum, feces, and saliva across all experimental groups on Day 35. (e) Long-term neutralizing activity of serially diluted fecal E18-3-specific IgG collected from the HBc-E18-3 floss group on Day 56. (f) Dynamic serially diluted serum anti-E18-3 IgG titers of the floss mucosal group versus the Freund’s adjuvant subcutaneous group at Day 0, 7, 21, and 35.

Day 56 detection revealed that fecal HBc-E18-3-specific IgG from the HBc-E18-3 floss group retained potent neutralizing activity even at a 1:12 dilution with minimal antibody decay, indicating durable long-term neutralizing capacity of the vaccine (Figure 3e). Serial serum dilution curves across Day 7, 21, and 35 demonstrated higher peak antibody titers in the subcutaneous injection group; however, floss immunization utilized only 25 μg antigen (vs. 100 μg for subcutaneous delivery) yet sustained stable long-term humoral responses (Figure 3f). Cross-group comparison across all sampling timepoints confirmed that mucosal secreted E18-3-specific IgA exhibited markedly weaker neutralizing activity than IgG.

Three experimental groups: HBc-CpG floss group, HBc-E18-3-CpG floss group, HBc-E18-3 Freund’s adjuvant subcutaneous group (n=6 mice per group). Data are presented as mean ± standard deviation; intergroup differences analyzed via Kruskal–Wallis test followed by Dunn’s post-hoc test, *P < 0.05. All plots show representative data from two independent replicate experiments.

### 3.4 Mechanistic Investigation of FcRn-Mediated IgG Responses Induced by VLPs and Favorable Systemic Safety Profile of Floss-Delivered HBc-E18-3

As shown in Figure 3d and 3e, antibodies generated via floss-mediated vaccination lacked neutralizing activity against isolated linear E18-3 peptide antigen. This observation suggests that cellular uptake, intracellular processing, and antigen presentation of exogenous antigens represent critical steps required to mount effective adaptive host immunity against most pathogens. A core rate-limiting step for antigen presentation is trafficking of exogenous cargo to intracellular compartments, where antigens are proteolytically processed and loaded onto MHC molecules. Loading of exogenous antigens onto MHC class II molecules and cross-presentation onto MHC class I molecules both occur within intracellular vesicular compartments. Accordingly, exogenous antigens require FcγR-directed internalization prior to entry into antigen-processing machinery, followed by FcRn-mediated intracellular shuttling of endocytosed antigen^34^.

Circulating IgG concentrations were markedly higher than IgA *in vivo*, a phenomenon attributed to FcRn-dependent IgG recycling that extends antibody serum half-life^35^. FcRn binds, transports, and recycles IgG to protect antibodies from lysosomal degradation, with vascular endothelial cells serving as the primary tissue compartment mediating FcRn-dependent IgG catabolic protection^36^. In autoimmune diseases driven by pathogenic autoantibodies, FcRn prolongs the half-life of autoreactive IgG and amplifies inflammatory signaling via active immune functions to exacerbate disease progression. In preclinical animal models including experimental autoimmune encephalomyelitis (EAE, a multiple sclerosis model), myasthenia gravis (MG), pemphigus, rheumatoid arthritis (RA), and inflammatory bowel disease (IBD), FcRn-knockout mice or animals treated with FcRn blocking agents displayed attenuated disease severity accompanied by reduced serum levels of pathogenic IgG. Given the central pathological role of FcRn in autoimmunity, blockade of FcRn–IgG interactions has emerged as a promising therapeutic strategy. The underlying mechanism relies on competitive inhibition to prevent recycling of pathogenic IgG (and total circulating IgG), accelerating lysosomal degradation and lowering serum antibody concentrations^35,37^.

Post-vaccination, serum IgG robustly neutralized antigens bearing native HBc conformational epitopes, while fecal IgG mediated specific antigen neutralization—findings consistent with FcRn-facilitated transcytosis of antigen-specific IgG across intestinal mucosal epithelia. Saliva from the HBc-CpG group neutralized HBc antigen, and saliva from the HBc-E18-3 Freund’s adjuvant group neutralized HBc-E18-3 antigen; in contrast, saliva collected from the HBc-E18-3-CpG floss group failed to exert antigen-specific neutralization, further corroborating this mechanistic model. FcRn is constitutively expressed across the human lifespan in antigen-presenting cells, intestinal epithelial cells, and select mucosal surfaces. It mediates bidirectional IgG transcytosis across intestinal epithelia to confer luminal immune protection and support mucosal immune surveillance^35^, increasing intestinal IgG permeability^38^. IgG forms immune complexes with luminal viral antigens to modulate adaptive immune responses against pathogens and foreign antigens^39^. FcRn captures luminal IgG–antigen immune complexes and shuttles them across epithelial monolayers to lamina propria dendritic cells, triggering robust CD4^⁺^ T cell activation and initiating pathogen-specific intestinal adaptive immunity.

Notably, fecal HBc-E18-3-specific IgG collected at Day 56 (1:12 dilution) yielded a 2-fold higher signal relative to HBc-specific IgG measured at Day 35 (1:2 dilution). This result implies that the linear epitope E18-3 derived from E18 VP1 localizes within the FcRn-binding interface of viral antigens, increasing binding affinity between HBc-E18-3-elicited IgG and FcRn and consequently enhancing FcRn-mediated IgG recycling and transepithelial transport efficiency.

Collectively, these mechanistic insights support a translational framework: peptide mimetics corresponding to epitopes targeted by pathogenic autoantibodies can be engineered into VLPs to generate competitive FcRn antagonists, continuously lowering circulating pathogenic IgG concentrations and preventing autoimmune disease relapse. All mice receiving floss-based mucosal vaccination exhibited normal physiological status with no hair loss, weight reduction, or decreased locomotor activity; mice receiving subcutaneous Freund’s adjuvant immunization displayed prominent dorsal alopecia and reduced mobility (Figure 4b). H&E staining of heart, liver, spleen, lung, and kidney tissue sections from all experimental groups revealed intact tissue architecture with no cellular degeneration, necrosis, massive inflammatory infiltration, congestion, or fibrotic lesions (Figure 4c), confirming minimal systemic visceral toxicity induced by floss-delivered mucosal VLPs.

**Figure 4.**
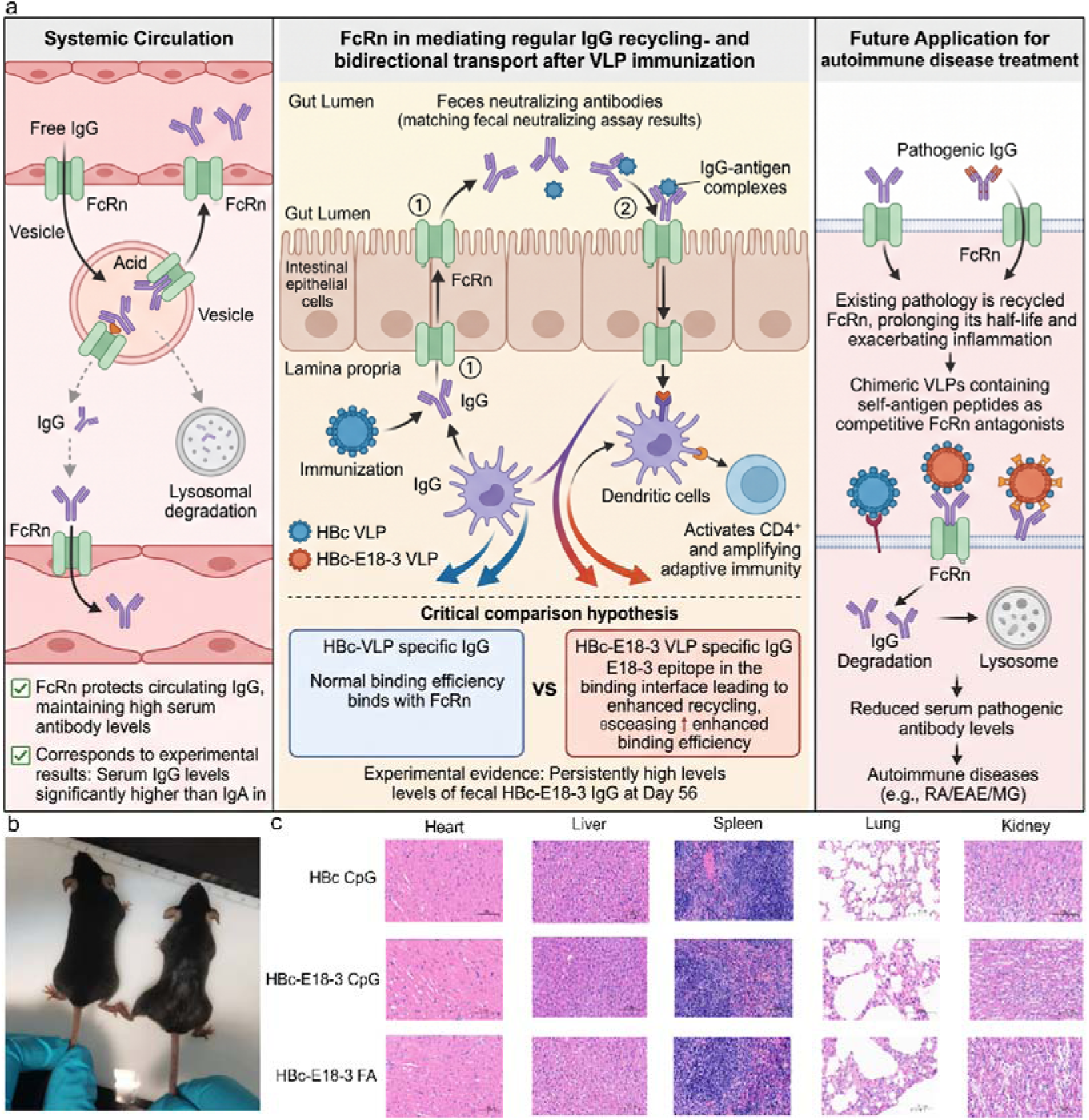
Speculation for FcRn-mediated circulatory protection and bidirectional mucosal transcytosis of VLP-elicited IgG, and prospective translational applications for autoimmune disease intervention. (a) Mechanistic schematic. Left panel: Vascular endothelial FcRn mediates recycling of circulating IgG to prevent lysosomal degradation and maintain high serum antibody concentrations. Middle intestinal epithelial compartment: FcRn orchestrates bidirectional IgG transcytosis. Antigen-specific IgG generated in the lamina propria is transported apically into the gut lumen to mediate fecal antigen neutralization; luminal IgG–antigen immune complexes are captured by FcRn and shuttled basolaterally to dendritic cells to activate CD4^⁺^ T cell-dependent adaptive immunity. A testable hypothesis is proposed based on experimental data: the E18-3 epitope resides within the FcRn-binding interface, conferring superior FcRn binding, recycling, and transcytosis efficiency to HBc-E18-3-specific IgG. Right translational panel: Chimeric VLPs embedded with autoantigen peptides act as competitive inhibitors blocking FcRn – pathogenic IgG interactions, accelerating lysosomal degradation of autoreactive antibodies as a potential intervention strategy for autoimmune disorders. (b) Representative photographs of experimental mice. (c) H&E-stained pathological sections of heart, liver, spleen, lung, and kidney tissues collected from all experimental groups.

## 4 Discussion

### 4.1 Translational Value of the Integrated In-House *evolution_conservation* Bioinformatic Pipeline

Traditional molecular evolutionary analysis of enteroviruses requires separate operation of independent software suites for NCBI batch retrieval, MAFFT alignment, MEGA-based phylogenetic reconstruction, and discrete Python statistical scripts, with repeated file import and export creating cumbersome workflows incompatible with rapid surveillance of emerging variant outbreaks. The automated *evolution_conservation* pipeline developed herein encapsulates the full analytical workflow within a single executable script, generating complete visualized phylogenetic outputs (Figure 1) following input of target sequence datasets. The resulting phylogenetic data enable rapid identification of genetically related historical viral strains, supporting retrospective evaluation of the safety and efficacy of prior prophylactic and therapeutic interventions to guide iterative vaccine optimization. This analytical tool is particularly valuable for rare post-E18 infectious sequelae including aseptic meningitis and poliomyelitis-like myasthenia gravis, facilitating global patient cohort localization.

### 4.2 Mechanistic Differences in Epitope Recognition Between Gingival Floss Immunization and Subcutaneous Injection

This study is the first to identify divergent epitope recognition and IgG neutralization profiles induced by subcutaneous versus floss-mediated gingival mucosal immunization using dual-epitope HBc-VLPs displaying both particulate HBc conformational epitopes and linear E18-3 peptides. The thin junctional epithelial barrier of the gingival sulcus efficiently internalizes nanoscale HBc particulate antigens, whereas isolated linear E18-3 peptides cannot be captured and processed by antigen-presenting cells, failing to elicit linear epitope-specific neutralizing antibodies. This disparity directly validates intact particulate HBc nanoparticle scaffolds as an indispensable delivery vehicle for non-invasive floss-based mucosal vaccination. Analogous immunological discrepancies have been documented in food allergy research, where oral antigen exposure triggers distinct immune responses relative to parenteral delivery^40^.

### 4.3 Dual Regulatory Axis of FcRn: Antiviral Prophylaxis and Autoimmune Disease Intervention Potential

ELISA results confirmed that floss-mediated gingival mucosal immunization induces sustained high concentrations of HBc-E18-3-specific IgG in both intestinal mucosal compartments and systemic circulation. Published literature confirms FcRn acts as an essential host receptor mediating E18 cellular entry and a central pathological driver of post-E18 autoimmune meningitis and generalized myasthenia gravis. Clinically approved anti-FcRn monoclonal antibodies such as efgartigimod and rozanolixizumab act via competitive occupancy of FcRn binding pockets to shorten the circulating half-life of pathogenic autoantibodies and alleviate neurological tissue injury, yet these therapeutics only provide transient symptomatic relief without primary blockade of viral infection or autoantibody generation^41^.

Drawing on therapeutic neoantigen vaccine paradigms developed to prevent tumor recurrence, the HBc-E18-3 mucosal VLP platform described herein establishes a novel framework for developing antigen-competitive prophylactic and therapeutic vaccines targeting FcRn-dependent myasthenia gravis and autoimmune encephalitis. Based on documented molecular interactions between IgG and FcRn, we propose that long-lived circulating E18-specific IgG competes with viral particles and autoreactive antibodies for epithelial FcRn occupancy to exert dual protective effects: (1) inhibition of E18 utilization of FcRn as an entry receptor to mediate direct viral neutralization; (2) suppression of pathogenic auto-IgG transcytosis across the blood–brain barrier to mitigate post-E18 autoimmune neurological damage.

Mucosal secreted IgA lacks FcRn-binding domains and only mediates transient luminal viral neutralization, while IgG persists long-term systemically via FcRn-dependent recycling to simultaneously execute antiviral neutralization and autoantibody blockade, forming a stratified multi-layered mucosal immune defense network.

### 4.4 Clinical Translational Advantages of Floss-Delivered Mucosal VLP Vaccines

Compared with conventional injectable vaccines, floss-mediated gingival mucosal delivery offers multiple practical translational benefits: low single-dose antigen dosage, simplified cold-chain storage requirements for protein-CpG adjuvant formulations, self-administration suitability for needle-phobic populations, and robust systemic safety validated via multi-organ pathological staining with no detectable visceral tissue damage—making this platform ideal for large-scale non-invasive mass vaccination campaigns. Truncated HBc scaffolds (residues 1–149) stably self-assemble into homogeneous T=4 nanoparticles free of exogenous genomic nucleic acid interference, serving as a universal modular display platform for diverse exogenous viral epitopes.

### 4.5 Study Limitations and Future Research Directions

1. The current iteration of the *evolution_conservation* pipeline is restricted to viral capsid protein sequences; future upgrades will incorporate whole-genome analytical modules. Constrained by laboratory resources, collaborative research with specialized structural biology groups will integrate cryo-electron microscopy and AI predictive modeling^42–45^ to resolve high-resolution three-dimensional structures of antigenic epitopes, IgG, and FcRn complexes and quantify competitive binding kinetics. Following neoantigen therapeutic vaccine design frameworks, subsequent engineering of HBc scaffolds will incorporate multiple target epitopes associated with FcRn-dependent autoimmune disorders to develop broad-spectrum mucosal therapeutic vaccines and expand non-invasive intervention strategies for neuroimmune diseases.
2. Mice are non-permissive natural hosts for E18 infection, limiting predictive translational value of viral challenge and autoimmune model intervention assays for human clinical outcomes, which were therefore not performed in this study. Comprehensive quantitative profiling of antigen-specific IgG and IgA across distinct immunization routes completed herein enables functional vaccine efficacy assessment via neutralization assays and mechanistic investigation of immune signaling pathways. Combined with phylogenetic patient cohort identification and AI-aided structural modeling, this bioinformatic workflow represents a regulatory-aligned in silico alternative to traditional animal experimentation.

In summary, this study constructed non-invasive floss-mediated gingival mucosal HBc-E18-3 VLPs using the pre-screened E18-3 epitope, paired with the custom-built integrated *evolution_conservation* bioinformatic pipeline for viral evolutionary profiling. This mucosal vaccine platform induces durable mucosal antigen-specific IgG for prophylaxis against circulating E18 strains, while establishing a reusable modular design paradigm for developing prophylactic and therapeutic vaccines targeting FcRn-mediated autoimmune meningitis and myasthenia gravis.

## Supporting information

S1 Full Python source code for the automated evolution_conservation analytical pipeline

## Acknowledgements

We sincerely thank Prof. Jie Wu of China Pharmaceutical University for laboratory infrastructure support for recombinant protein production, VLP characterization, mouse gingival mucosal immunization, and pathological tissue analysis. We acknowledge all members of the Wu laboratory for daily technical support and academic discussion facilitating successful completion of all experimental work.

## Ethics Statement

All animal experimental protocols were approved by the Animal Ethics Committee of China Pharmaceutical University, and all animal handling procedures strictly complied with the internationally recognized 3R principles for laboratory animal welfare.

## Supplementary Materials List

S1 Full Python source code for the automated *evolution_conservation* analytical pipeline (complete scripts for phylogenetic tree generation and downstream quantitative analysis)

S2 Multiple sequence alignment results of E18 VP1 amino acid sequences

a Filtered_unique_vp1(Alignment of non-redundant Echovirus VP1 sequences following quality control filtering)

b Mafft_alignment(Sequence conservation visualization generated from high-accuracy global MAFFT alignment, with asterisks marking fully conserved amino acid residues)

